# Highly sensitive enzyme- and amplification-free, quantitative DNA detection using YVO_4_:Eu luminescent nanoparticle probes

**DOI:** 10.64898/2026.02.24.707628

**Authors:** Robin Kuhner, Christophe Cardone, Rafael Vieira Perrella, Rabei Mohammedi, Thierry Gacoin, Roxane Lestini, Cedric.I Bouzigues, Antigoni Alexandrou

**Affiliations:** Laboratoire d’Optique et Biosciences, CNRS, INSERM, Ecole Polytechnique, Institut Polytechnique de Paris, Palaiseau, France; Laboratoire de Physique de la Matière Condensée, CNRS, Ecole Polytechnique, Institut Polytechnique de Paris, Palaiseau, France

## Abstract

The sensitive detection of nucleic acids is crucial for the accurate diagnosis of infections. In this context, amplification-based methods, such as the quantitative Polymerase Chain Reaction (qPCR) are the gold standard for ultrasensitive DNA or RNA detection and quantification. However, despite its widespread use in developed countries during the COVID-19 pandemic, qPCR remains a costly tool, difficult to implement into low-infrastructure locations. Efforts for the development of alternative tools have yielded high sensitivity approaches but sensitivity is typically reached at the expense of complexity. We here report the development of a simple, sensitive, amplification-, and enzyme-free nucleic acid detection technique using YVO_4_:Eu luminescent nanoparticles. We established an optimized interaction scheme to efficiently reveal target DNA fragments with nanoparticles. By exploiting the extremely strong absorption of the vanadate matrix in the UV to excite the nanoparticles inducing the characteristic Eu^3+^ emission at 617 nm via energy transfer, we achieved a highly sensitive (down to 500 particles/mm^2^; 17,000 particles/well) read-out in standard microplates using a home-made optical reader with light-emitting diode (LED), 275-nm excitation. We reached a 50-aM (30,000 copies/mL) sensitivity for the detection of the 72-base DNA fragment of the SARS-CoV-2 *n1* gene. Our new quantitative analytical method detects nucleic acids without amplification with performances close to standard PCR (10,000 copies/mL)^1^, and could be the basis for a transportable alternative for the diagnosis of infectious diseases.

---

The detection of pathogen (bacterium, virus or parasite) DNA/RNA is the reference diagnosis technique in many cases.^2–4^ Moreover, circulating nucleic acids^5,6^ are increasingly used for diagnostic purposes. Today, a variety of ultrasensitive techniques is available for nucleic acid detection: polymerase chain reaction (PCR), reverse transcription PCR (RT-PCR, for the detection of RNA), quantitative PCR (qPCR),^7^ or loop-mediated isothermal amplification (LAMP)^8^.

However, these methods require amplification of the nucleic acid – typically 30 to 40 cycles - and involve the use of costly enzymes. Moreover, the most sensitive of these techniques, qPCR, also requires expensive equipment, trained personnel and sample transfer to the equipment location. In low-infrastructure locations, such equipment and trained personnel are highly scarce. Additionally, polymerization enzymes may be inhibited by components present in unpurified or environmental samples. Concerning the detection of short circulating nucleic acids, design of appropriate primers may be highly complex.

Standard PCR and LAMP are less costly and in particular LAMP is compatible with low-resource settings, but both are less sensitive and do not provide quantitative results. Standard PCR apparatus typically detects DNA concentrations down to 20 aM, *i. e*. approximately 10,000 molecules/mL, about 2 orders of magnitude less sensitive than qPCR which detects down to about 10-100 molecules/mL, 0.1 aM or less^7,9^. These orders of magnitude are indicative and depend on the type of detected nucleic acid.

To quantitatively measure nucleic acid concentrations, so-called PCR-ELISA approaches have been proposed^10,11^. Such techniques, however, still rely on amplification reactions and involve a two-step procedure. More recently, recombinase polymerase amplification was combined with horseradish peroxidase (HRP) or surface-enhanced Raman scattering (SERS) detection^12^ and RNA aptaswitches with LAMP to reach 1-2 aM detection.^13^ Alternatively, molecular biology tools like toehold entropy-driven chain displacement reactions have been exploited to produce large quantities of a secondary DNA strand that is subsequently detected in an enzyme-free amplification scheme^14,15^. This can then be combined with the opening of hairpin RNA structures using an endonuclease^15^ or by molecular-beacon mediated isothermal circular strand displacement polymerization reaction^14^ to reach limits of detection down to 7.8 aM, and 6 aM, respectively.

Alternative approaches have been proposed aiming at improving sensitivity through signal enhancement using fluorescence or electroluminescence detection. For instance, silver nanoparticle aggregation in the presence of target DNA enabled detection down to 50 fM on fluorescent microarrays.^16^ A mixture of gold nanotriangles and nanoparticles, together with [Ru(bpy)_3_]^2^ and carbon nanodots as coreactants, provided a nanostructured electroluminescent platform with detection limits down to 514 aM.^17^ More elaborate signal enhancement approaches resort to macrocyclic host molecules, calixarenes, for enrichment of the electrochemical mediator, to form concatamers of these macromolecules reaching 3-aM detection of the SARS-CoV-2 ORF1ab^18^.

These alternative approaches do not rely on DNA amplification, but the achieved higher sensitivity still falls short of PCR performances and/or comes at the expense of complexity. We here propose a sensitive, quantitative nucleic acid detection method based on a simple sandwich reactional scheme, where complementary oligonucleotides are the capture and detection recognition elements (Fig. 1) and ultrabright, photostable, luminescent rare-earth-based nanoparticles are used as probes.

**Fig. 1.**
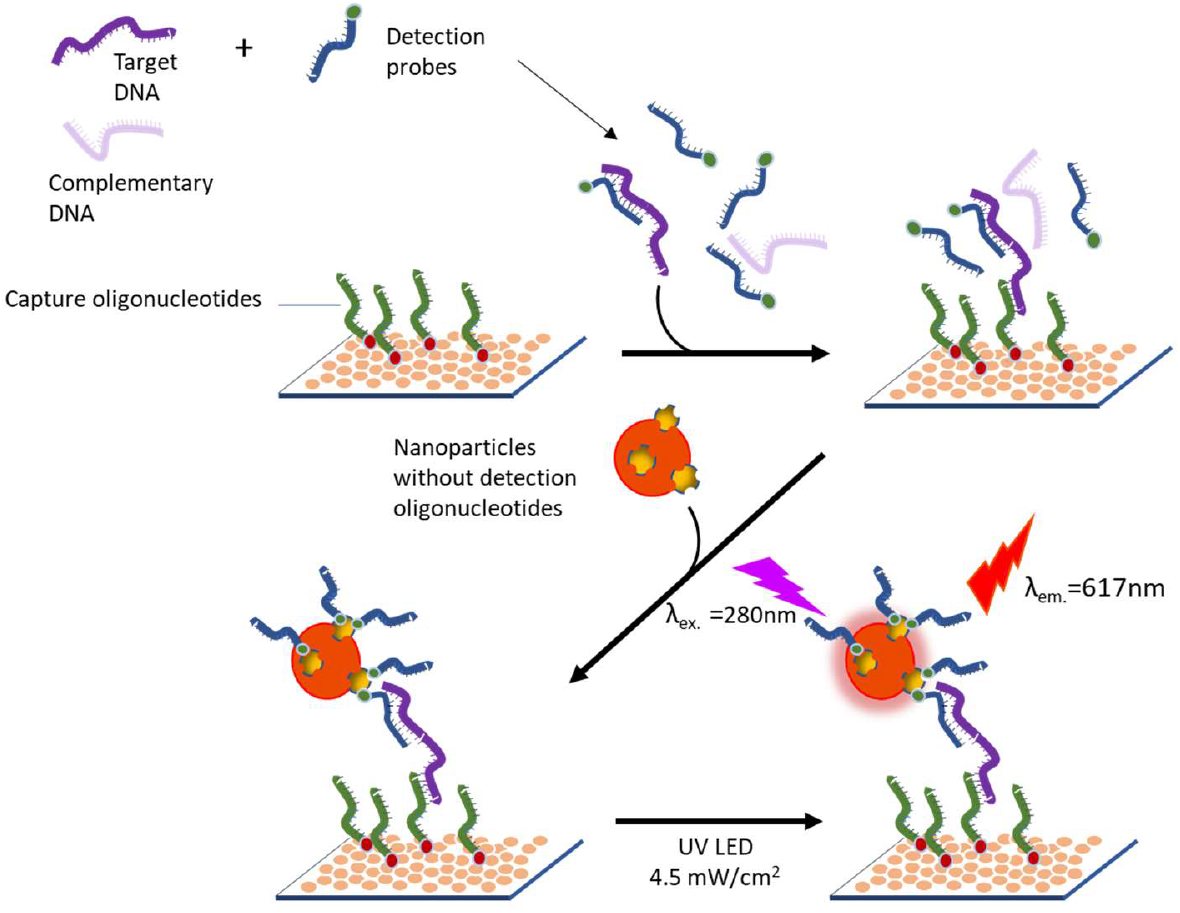
A) DNA detection with capture and detection oligonucleotides using YVO_4_:Eu nanoparticles as probes: in denaturing conditions, the target DNA is simultaneously incubated with biotin-labeled detection oligonucleotides and with the capture oligonucleotide-coated surface (top right). YVO_4_:Eu nanoparticles coupled to streptavidin are then incubated with the target DNA captured on the surface (bottom left). After rinsing, surface-bound nanoparticle probes are excited at 275 nm and their emitted light is detected (bottom right).

We have recently shown that yttrium vanadate 38-nm nanoparticles doped with Eu^3+^ ions used as probes enable ultrasensitive femtomolar detection of proteins.^19^ The analyte detection benefits from the ultrastrong absorption extinction coefficient of the vanadate matrix at 280 nm, 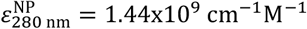, and is based on a home-made transportable microplate reader (Supporting material).^19^ Further advantages of these Eu-doped nanoparticles are : i) their narrow emission and large Stokes shift which enables the use a narrow filters for the rejection of parasitic emissions, ii) their high photostability and absence of blinking, and iii) facile synthesis and functionalization. 20-mer oligonucleotides typically show higher affinity with their DNA targets than antibodies for their antigens;^20^ we therefore expect to reach sub-fM sensitivity.

We synthesized YVO_4_:Eu nanoparticles exploiting a metavanadate (VO_3_^-^) instead of an orthovanadate ion (VO_4_^3-^) precursor together with three equivalents of NaOH which yields higher reproducibility and crystallinity (Supporting material) ^21^. The quantum yield was then enhanced by a post-synthesis heating step at 200 °C to reach 18%. The obtained 38-nm YVO_4_:Eu (20%) nanoparticles (Fig. S1a) were then functionalized using previously published protocols^22–24^ and coupled to streptavidin molecules to yield the detection probes that recognize biotinylated detection oligonucleotides, complementary to the target strand. These nanoparticles absorb at 280 nm (absorption of the vanadate groups) and, after energy transfer to Eu^3+^ ions, show a main emission peak at 617 nm (Fig. S1b).^19^

The *in situ* optical performance of the nanoparticles, *i. e*. the detection sensitivity expressed in nanoparticle surface density, was determined as follows. We non-specifically adsorbed nanoparticles on microwell plates by incubation with nanoparticle solutions. We then determined for samples of different concentrations the numbers of surface-bound single nanoparticles detected through a wide-field epifluorescence microscope (Fig. 2A-B) that were compared to the photomultiplier (PMT) signals recorded by the home-made reader for the same sample. Multiplying by the ratio of the well surface (well diameter of 6.5 mm) to that of the microscope field-of-view (142.3 × 142.3 μm^2^) yields the total number of surface-bound nanoparticles in a well, which leads to a calibration of the reader output voltage in absolute nanoparticle number and density (Fig. 2C). The minimal detectable nanoparticle signal – detectable above the blank signal background due to non-specific interactions and its fluctuations - was 10 mV, corresponding to a surface density of 500 NP/mm^2^. This corresponds to only a few nanoparticles detected per field-of-view (Fig. 2A) and means that our ensemble nanoparticle detection reader is as sensitive as a high-end microscope operating in the single-molecule regime. Moreover, in the investigated range (1100-177,000 NP/mm^2^), the PMT signal is a linear function of the adsorbed nanoparticle number, which enables a direct read-out of the nanoparticle number.^19^

**Fig. 2.**
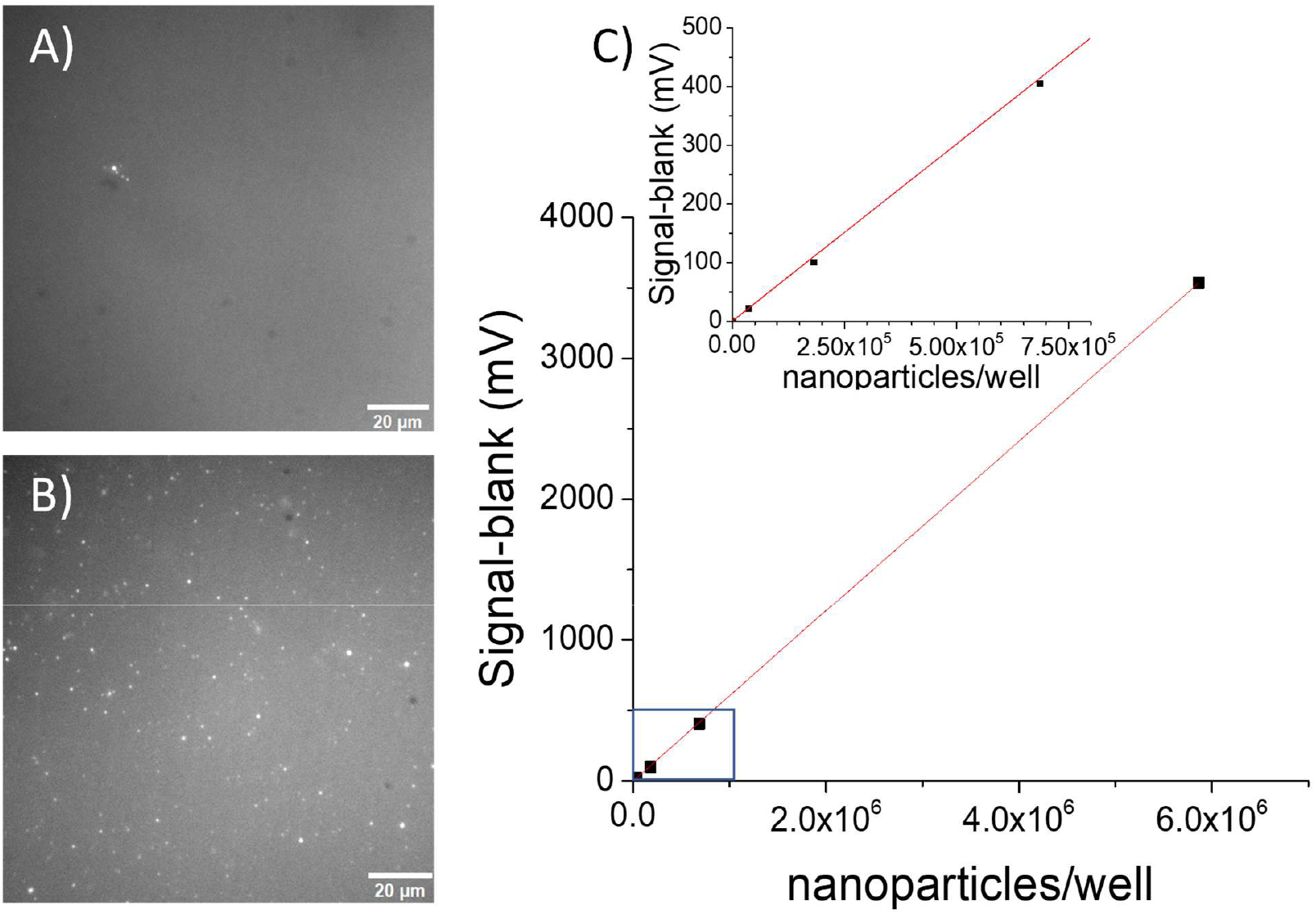
A-B) Wide-field microscopy images of microplate wells yielding low and intermediate signal values with the microplate reader of 22 mV A) and 415 mV B), respectively. Very few particles are observable in (A), whereas tens of particles are seen in B). Acquisition time: 500 ms. C) Calibration curve relating the number of nanoparticles per microscope field of view (converted to particle number/well) to the PMT signal obtained with the plate reader and linear fit of the data (red line). The blank signal (155 mV) was subtracted from the signal values. Inset: zoom-in corresponding to the blue rectangle.

In a classical sandwich ELISA for protein detection, a capture antibody is deposited at the bottom of the well. The target molecule or analyte is then added (step 1) whose presence is detected using enzyme-labeled detection antibodies bound to a probe (step 2). Each reaction step is followed by rinsing to remove unbound molecules (four steps in total). We propose a modified transposition of this approach to the detection of nucleic acids (Fig. 1): (i) the target nucleic acid is incubated with detection oligonucleotides (ON) in large excess with the surface coated with capture ON, also in large excess (45 minutes) (ii) streptavidin-coated nanoparticles attach to the biotinylated surface-bound detection ON (1 hour). We ensure that both the total number of streptavidin molecules coupled to the surface of the nanoparticles and the total number of particles is in excess with respect to the total number of detection oligonucleotides (ratios of 180 and 4.5, respectively; Table S1). NaOH is added in the reaction solution of step (i) either to dehybridize double-stranded target DNA (dsDNA) or to remove potential secondary structure of single-stranded target DNA (ssDNA). After a single rinsing step (total of three steps), the reader measures the bound particle luminescence. A rinsing step after the incubation of the target DNA with detection and surface-bound capture ON is unnecessary. Any free nanoparticle-detection ON complexes in solution are removed during the final rinsing step thus minimizes the total number of steps. This protocol aims at DNA interactions (in the whole concentration range of interest) with capture and detection ON in large excess (Table S1) resulting in binding to almost the entire population of target molecules.

A direct transposition of the ELISA protocol with detection ON extemporaneously labelled by nanoparticles lead to poor sensitivity (data not shown) probably due to the large size of the nanoparticles which may slow down the hybridization kinetics, and reduce the labelling efficiency. Notably in the case of dsDNA detection, nanoparticle-labelled detection ON are in competition with the native complementary DNA strand. Therefore, even when in large excess, we expect these nanoparticle-labelled ON to diffuse more slowly in solution than unlabelled detection ON, too slowly to preferentially bind to the target DNA.

The capture surface is coated with capture ON complementary to the target strand as follows: First, anti-DIG antibodies are adsorbed at the bottom of a 96-well plate (Fig. 1). After washing and passivation, the DIG-labeled capture oligonucleotide is added in large excess, *i. e*. five times more molecules than the initial number of incubated antibody molecules (Table S1).

We detected the 72-bp *n1* gene of the SARS-CoV-2 virus in three different forms: i) dsDNA obtained through PCR with the appropriate primers on the *n1*-containing plasmid pEX-A128-nCOV N1 (PCR*n1*), ii) chemically synthesized ssDNA (ss *n1*), iii) and dsDNA obtained by hybridization of the two complementary synthesized single strains (ds *n1*) (Fig. 3). The PCR products were characterized with electrophoresis: three bands were observed corresponding to the *n1*-gene PCR fragment at the expected molecular weight (MW) and two remains of the released plasmid and the supercoiled plasmid, respectively (Fig. 3A). Image analysis shows that the short PCR fragment contributes to 70 ± 11 % of the total fluorescence signal corresponding to 99.5% of the DNA fragments), which ensures a high level of purity of the sample before detection.

**Fig. 3.**
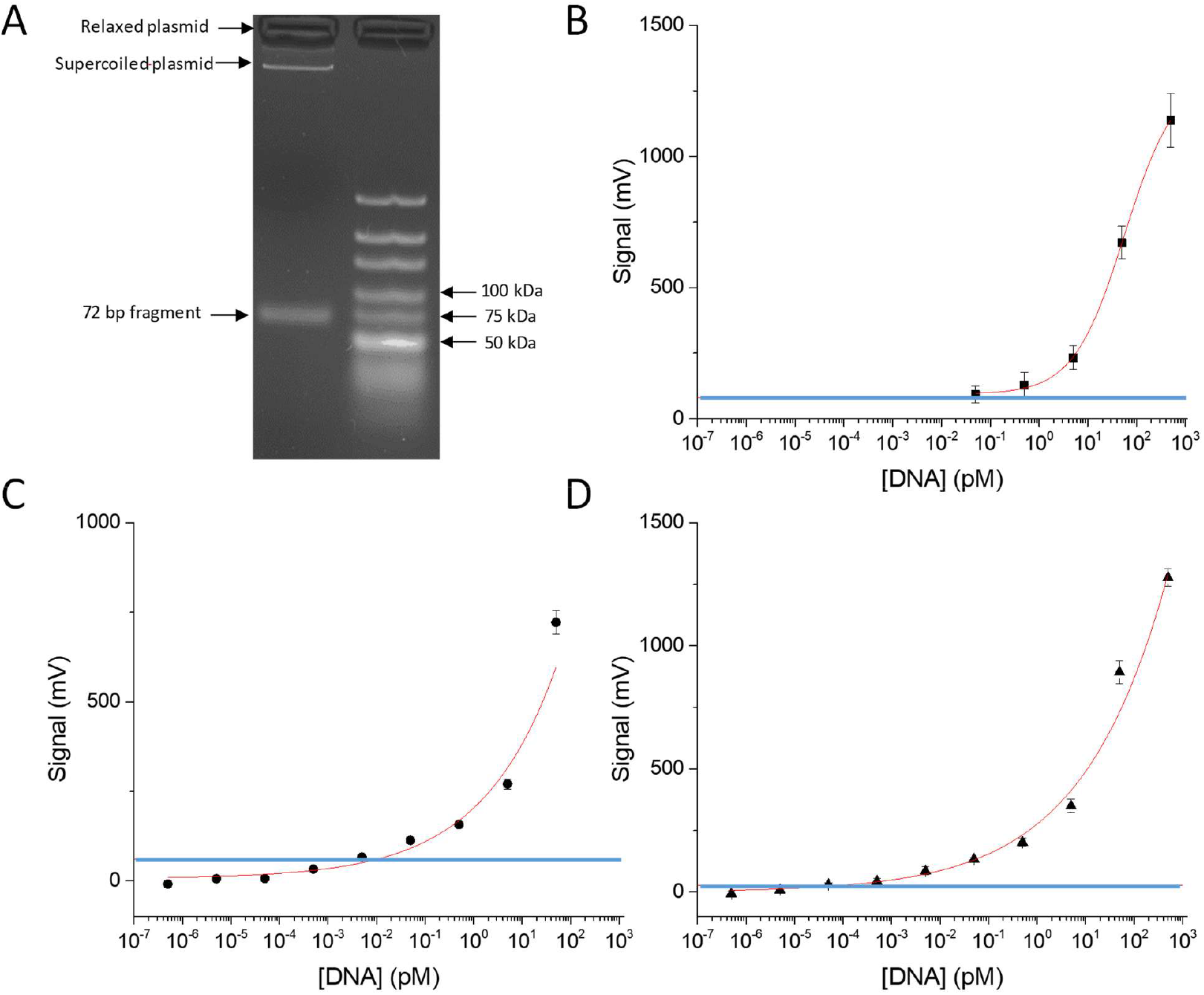
Detection of different configurations of the n1 gene of the SARS-CoV-2 virus in buffer. A) Electrophoresis of the PCR product in agarose gel 4% (left) together with an ultra-low MW ruler (right). B) Detection of the ds n1 gene obtained through PCR amplification of an n1-gene containing plasmid. Detection of a synthetized dsDNA C) and ssDNA D) of the 72-nucleotide n1 gene. The data have been fitted with the Hill equation: n= 0.83±0.07, 0.28±0.14, and 0.25±0.06 in B), C), and D), respectively. Blue horizontal lines correspond to three standard deviations of the blank signal. The values of the average blank signal, 255.3, 237.8 mV, and 201.9 mV (n=10 in each case) in B), C), and D), respectively, have been subtracted from the average signal values. Experiments performed in triplicate. Error bars show the standard deviation.

All fragments could be quantitatively detected over an extended concentration range (at least between 1 and 1000 pM; Fig. 3). We determined the signal corresponding to the limit of detection (LOD), defined as Signal(*LOD*) = Signal(0) + 3*σ*_0_, where Signal(0) and *σ*_0_ are the blank signal and its standard deviation, respectively (blue lines in Fig. 3). The lowest measured concentration above this signal was: *LOD*_*PCRn*1_ = 50 fM, *LOD*_*ds n*1_ = 5 fM, and *LOD*_*ss n*1_ = 50 aM (the Hill equation fits cross the LOD signal line at 10 fM and 109 aM for *ds n1* and *ss n1*, respectively). The difference in LOD for the PCR fragment and the synthesized dsDNA can be explained by the lower purity of the PCR product. Even though the dsDNA target represents more than 99% of the molecules in this sample, the presence of bulky plasmid material may decrease the interaction probability between the target DNA and the detection and capture oligonucleotides by introducing steric constraints to the diffusion process. A further decrease of the LOD for the synthesized ssDNA compared to the ds one yields an LOD of only 30 000 copies/mL (given the sample volume of 100 μL) without any enzyme-dependent or other amplification step. In the ssDNA case, there is no competition between the re-hybridation of the target complementary strands and the association between target fragment and detection or capture ON.

Although these measurements were performed in controlled buffers and should be reproduced in native biological samples, the presence of a high concentration of salmon sperm DNA in the sample, for passivation reasons, ensures the specificity of our test. The non-specific signal in the presence of this additional DNA remains indeed limited, indicating a high sensitivity/specificity detection of the target DNA. This highlights the potential of our approach for efficient detection in complex samples, in which (q)PCR sensitivity is notably degraded.

In conclusion, thanks to the unique optical properties of YVO_4_:Eu nanoparticle probes, we obtained amplification- and enzyme-free yet sensitive attomolar DNA detection with a straightforward scheme. It requires only an inexpensive reader mainly consisting of a 100-mW LED emitting at 275 nm and a photomultiplier tube. Moreover, our approach is inherently multiplexable because it employs multiwell microplates. This new method offers the advantage of achieving detection limits down to 50 aM, with similar sensitivity as standard PCR, without relying on complex and expensive macromolecules or requiring enzymatic amplification at any stage of the protocol. This direct sandwich double hybrization approach should thus provide an inexpensive alternative to current polymerase-chain-amplification approaches, which are both impaired by the presence of other biomolecules in non-purified samples and not accessible in low-infrastructure environments.^14,15^

## Supporting information

Materials and methods, Fig. S1, Table S1.

## Acknowledgments

We gratefully acknowledge financial support from the Centre Interdisciplinaire d’Études pour la Défense et la Sécurité (CIEDS) under the DAVID and DOMUS projects.

