## supporting information for "Highly sensitive enzyme- and amplification-free, quantitative DNA detection using YVO_4_:Eu luminescent nanoparticle probes"


**Nanoparticle size and excitation/emission spectra**

The nanoparticle size was determined from the analysis of individual particles (N=262) in TEM images Fig. S1(a). The average value of the short and long axis was determined to be 31.7 and 43.2 nm, respectively. By considering the third axis to be equal to the short axis, we calculated the mean particle volume to be 28 900 nm$^3$. The corresponding size for spherical particles with the same volume is 38 nm. For further details, see Ref. [1]

The excitation and emission spectra of the nanoparticles are shown in [Fig. S1(b)].

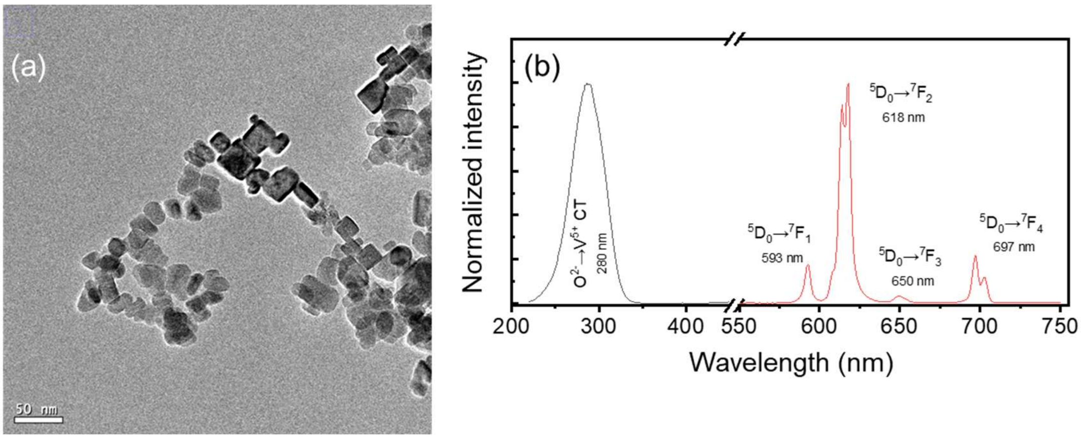


***Figure S1.*** *(a) High-resolution TEM image (scale bar: 50 nm) and (c) Excitation ($\lambda_{em}$ = 618 nm, black line) and emission ($\lambda_{exc}$ = 280 nm, red line) spectra of the colloidal suspension of silicated nanoparticles (~50 mmol.L$^{-1}$ in vanadate ions). The labels correspond to the $Eu^{3+}$ $^5D_0\rightarrow{}^7F_J$ (J=1-4) transitions (emission) and $O^{2-}\rightarrow V^{5+}$ charge transfer band (CT, excitation).*

| | | | |
|---|---|---|---|
| Surface preparation | Concentration of capture DIG-labeled oligonucleotides (nM) | Concentration of anti-DIG antibodies (nM) | Ratio anti-DIG antibodies/capture DIG-labeled oligonucleotides |
| | $3.33 \times 10^5$ | $6.67 \times 10^4$ | 5 |
| Step 1 | Concentration of detection biotin-labeled oligonucleotides (nM) | DNA target concentration (nM) | Ratio detection biotin-labeled oligonucleotides/DNA target |
| | 1 | 0.001 | 1000 |
| Step 1 | Concentration of capture DIG-labeled oligonucleotides (nM) | DNA target concentration (nM) | Ratio capture DIG-labeled oligonucleotides/DNA target |
| | $3.33 \times 10^5$ | 0.001 | $3.33 \times 10^8$ |
| Step 2 | Streptavidin concentration (conjugated to nanoparticles) (nM) | Concentration of detection biotin-labeled oligonucleotides (nM) | Ratio streptavidin molecules/detection biotin-labeled oligonucleotides |
| | 180 | 1 | 180 |
| Step 2 | Nanoparticle concentration (nM) | Concentration of detection biotin-labeled oligonucleotides (nM) | Ratio nanoparticles/detection biotin-labeled oligonucleotides |
| | 4.5 | 1 | 4.5 |

***Table S1.*** *Concentrations and excess ratios between the different reactants involved for a DNA target concentration of 1 pM. The left column indicates the step of the DNA reaction protocol.*

### Materials and methods

#### Chemicals

$Y(NO_3)_3.6H_2O$ (99.9%), $Eu(NO_3)_3.5H_2O$ (99.9%), sodium metavanadate ($NaVO_3$, 98%+), and (3-aminopropyl)triethoxysilane (APTES, 99%) were provided by Sigma-Aldrich and used without further purification. Sodium silicate solution ($Na_2O(SiO_2)_X.xH_2O$, extra pure, ~26.5% $SiO_2$ w/w, Merck), sodium hydroxide (1 mol $L^{-1}$ NaOH, Merck), anhydrous ethanol (99.9%, Carlo Elba), and N,N-dimethylformamide (DMF, 99.9%, Carlo Elba) were also used as received from Sigma-Aldrich anhydrous succinic (≥99%) , 1-ethyl-3-(3-dimethylaminopropyl) carbodiimide (EDC ≥90%, Sigma) and N-hydroxysulfosuccinimide (Sulfo-NHS ≥98%), Ethanol absolute anhydrous (99.9%), sodium chloride (NaCl, ≥99%). Tween 20 (≥40 %) potassium hydroxide (KOH, pellets, ≥86%, Fluka), mPEG-$NH_2$ (MW 500 g $mol^{-1}$, ≥95%, Biochempeg), anti-digoxigenin antibody (anti-Dig, Abcam), bovine serum albumin (BSA, ≥95%, Fischer Scientific) sodium dodecyl sulphate solution (SDS, 10 wt% in $H_2O$,) -(N-morpholino)ethanesulfonic acid (MES, ≥99%), streptavidin (from *Streptomyces avidinii*, essentially salt-free, lyophilized powder, ≥ 13 u/mg protein) N-lauroylsarcosine sodium salt (≥94%, Merck)

#### Synthesis of $YVO_4:Eu^{3+}$ nanoparticles

$Eu^{3+}$-doped $YVO_4$ nanoparticles (20 mol% of $Eu^{3+}$ doping) were prepared following the co-precipitation method described in our previous work.[2] Briefly, 25 mL of a 0.1 mol.$L^{-1}$ $NaVO_3$ solution containing 3 equivalents of NaOH with respect to $VO_3^-$ were freshly prepared (*i.e.* 0.3048 g of $NaVO_3$ dissolved in 17.5 mL of Milli-Q water and 7.5 mL of 1 mol.$L^{-1}$ NaOH solution). Then, 25 mL of a 0.1 mol $L^{-1}$ aqueous solution containing $Y(NO_3)_3$ (0.08 mol.$L^{-1}$) and $Eu(NO_3)_3$ (0.02 mol.$L^{-1}$) were added into the vanadate solution and left under vigorous stirring for 12 days at room temperature. The resulting suspension was centrifuged at 26323*g* for 30 min, and the pellet was resuspended in 50 mL of Milli-Q water. These centrifugation/redispersion steps were repeated two times to ensure a final conductivity < 100 µS.$cm^{-1}$ for the colloidal suspension.

Particles were then coated with silicate groups to avoid aggregation during subsequent annealing. In a typical procedure, 4.1 mL of a $Na_2O(SiO_2)_X.xH_2O$ solution were added dropwise to 50 mL of the particle suspension (50 mmol.$L^{-1}$ in vanadate). After 2 h under stirring, the particles were purified as detailed above. The resulting suspension, with a volume adjusted to 50 mL, was then centrifuged at 100*g* for 5 min to remove possible aggregates. Afterwards, silicate-coated nanoparticles were annealed at 200 °C for 2 h in a microwave (CEM Discover SP + Explorer). Finally, the solids were re-silicated as previously described to reinforce their colloidal stability. The obtained suspension of $YVO_4:Eu^{3+}$ nanoparticles (50 $mmol.L^{-1}$ in vanadate ions) was homogeneous and slightly light diffusing.

**Nanoparticle characterization**

Transmission electron microscopy was performed on a JEOL JEM-2010F transmission electron microscope (HT objective lens: Cs = 1.4 mm, Cc = 1.8 mm, Focal length = 2.7 mm) operating at 200 kV accelerating voltage. Prior to the observation, samples were prepared by depositing 20 µL of particle suspensions on a Parafilm. The carbon-coated 200 mesh copper grid, previously treated with a positive glow-discharge, was put in contact with the sample droplet (50 $mmol.L^{-1}$ in vanadate ions) for ~1 min. Then, the grid was rapidly removed and washed (~30 s) sequentially with two droplets of Milli-Q water. The fluid in excess was then carefully removed by placing filter paper onto the edge of the grid, followed by air drying. The observations were carried out at 20,000x, 80,000x, 200,000x, and 400,000x magnifications for different defocus values (defocus range = -0.2 to -1.5 µm) using a Gatan US4000 CCD camera.

Dynamic light scattering (DLS) and ζ-potential profiles were obtained on a Malvern Zetasizer Nano ZS using 0.1 $mg.mL^{-1}$ dispersions in Milli-Q water. Luminescence and excitation spectra were acquired on a Hitachi F-4500 spectrofluorometer. The emission quantum yields of colloidal suspensions (~50 $\mu mol.L^{-1}$ in vanadate ions) were estimated by comparing their integrated emission intensity (from the $^5D_0$ state of $Eu^{3+}$ ions) with the emission from a Rhodamine 6G solution ($10^{-5}$ mol $L^{-1}$ in ethanol, $q_{ST} = 0.94$)[3] having the same optical density (OD ~ 0.2) and excited at the same wavelength (280 nm). The vanadate concentration, $[VO_4^{3-}]$, was determined from absorbance measurements on dissolved particles following the method described by Kuhner et al.[1] and using the relation: $[VO_4^{3-}] = A_{280} / \varepsilon_{280}$, where $\varepsilon_{280}$ $(VO_4^{3-})$ = 4000 $M^{-1}$ $cm^{-1}$.[4] The nanoparticle concentration was then determined from the vanadate concentration, the average particle volume (see above), the volume of the crystalline unit cell (0.32 $nm^3$) [5]and the number of vanadate ions per unit cell (4).

**Nanoparticle functionnalization**

Nanoparticles were functionalized according to previous work[3] to enable their covalent coupling with oligonucleotides. To this end, 75 mL of re-silicated $YVO_4:Eu^{3+}$ nanoparticles at 3 mM in vanadate ions were added dropwise (1 $mL.min^{-1}$) to 225 mL of boiling ethanol containing 265 µL of APTES. The solution was left under reflux for 24 hours at 92 °C to ensure a uniform cross-linked aminated network surrounding the particles. The solids were then separated by centrifugation (26323*g* for 30 min), washed two times using an ethanol/water mixture (3/1 v/v), and redispersed in 5 mL of Milli-Q water. Finally, the colloidal suspension was added dropwise to 20 mL of anhydrous succinic in DMF (0.25 g $mL^{-1}$) and stirred overnight at room temperature. The particles were purified by 3 cycles of centrifugation/redispersion in the ethanol/water mixture and resuspended in 5 mL of 2-(N-morpholino)ethane sulfonic acid (MES) buffer (50 mM, pH 5 to 6). The different functionalization steps were characterized using ζ-potential measurements as discussed in Ref. [1]

**Bioconjugation to streptavidin**

150 µL of –COOH coated nanoparticles at a concentration of 200 nM were centrifuged for 15 minutes at 13700*g* and the supernatant was removed. 1-ethyl-3-(3-dimethylaminopropyl) carbodiimide (EDC) and N-hydroxysulfosuccinimide (Sulfo-NHS) were solubilized at a concentration of 50 mg/mL each in a 50 mM MES buffer at pH 5.5. This solution was quickly added to the nanoparticle pellet and the sample was sonicated for 15 seconds at 50% amplitude (Ultrasonic processor GEX130, tip ref. 423-A). After incubation for 25 minutes at 25°C under stirring at 300 rpm (Eppendorf Thermomixer C), the sample was centrifuged for 15 minutes at 15000*g* and the nanoparticle pellet was redispersed in 50 mM phosphate buffer at pH 7.4 by sonication during 15 seconds at 50 % amplitude. 40 equivalents of streptavidin molecules (Sigma, s4762-10MG) with respect to the nanoparticles were added and the sample was incubated at 25 °C with agitation at 800 rpm for 2.5 hours. The sample was centrifuged for 15 minutes at 13700*g* and the pellet was resuspended in 500 µL of blocking buffer (50 mM phosphate buffer at pH 7.4 containing 2% wt mPEG-$NH_2$ 500 (MF001005-500 Biochempeg) by pulsed sonication (1 s on/1 s off for 10 s at 20% amplitude) on ice. Then, the sample was incubated at 25°C for 1 hour with stirring at 300 rpm (Eppendorf Thermomixer C) and centrifuged for 15 minutes at 13700*g*. Finally, the resultant pellet corresponding to the streptavidin-nanoparticle conjugates was redispersed in conservation buffer (Tris 20 mM, pH 8, 1% wt BSA) and stored at -80°C.

**Optical reader**

The home-made reader integrates a 100-mW, 275-nm light-emitting diode (LED; Hex-S6060-DR250-W275-P100-V6.5, Laser Components) for excitation of the nanoparticle vanadate matrix, a 1-mm diaphragm restricting the illumination area to prevent excitation of multiple adjacent wells of the microplate, a photomultiplier tube (PMT, PMM02, Thorlabs), and two stepper motors (17HS15-1504S-X1), a lead screw in the Y direction and a pulley-belt system in the X direction, for microplate (655097, Greiner) readout. Emitted luminescence from immobilized nanoparticles in a microplate well is collected through a lens (0.79 numerical aperture, ACL25416U-A, Thorlabs), and focused onto the PMT with a f=100 mm lens (LA1509-A, Thorlabs). The PMT is not sensitive to UV light and is fitted with two interference filters selecting the nanoparticle luminescence centered at 617 nm (FF01-620/14-25, Semrock). This optical configuration suppresses most of the parasitic light coming from the excitation source or from the fluorescence of molecules and materials in the sample or in the reader. Control measurements confirmed that there is no crosstalk between different wells and that the signal across the microplate is homogeneous (for further details see Ref. [1]).

**Optical microscopy**

Suspensions of streptavidin-conjugated $YVO_4:Eu^{3+}$ nanoparticles were prepared at several concentrations in PBS buffer at pH 7.4. Aliquots of 100 µL were dispensed in duplicate into multiwell plates with a 190-µm thick cyclo-olefin bottom (Greiner Screenstar, Dutscher 655866). The plate was incubated for 48 hours at 4 °C under stirring at 300 rpm (Eppendorf Thermomixer C) followed by five washing steps with PBS buffer at pH 7.4 before reading on the homemade UV microplate reader to determine the signal in mV. Then each well was observed under a wide-field microscope for single-particle observation (Olympus IX83 equipped with a 63x plan-apochromat Zeiss objective, Ref. 44 07 62) and a Photometrics Evolve EM-CCD 512x512 camera) using excitation with a 466-nm diode laser (Modulight). Five independent regions of interest were recorded per well and analysed with a homemade MATLAB script to detect, count, and average the number of single nanoparticles visible in each field-of-view (142.3 µm x 142.3 µm).

**Nucleic acid detection**

Anti-Digoxigenin (DIG) antibody (Ab64509, Abcam) was applied to the bottom of multiwell plates at 10 µg/mL in 100 µL of PBS buffer pH 7.2 and incubated overnight at 4°C under stirring. The plate was then washed three times with 200 µL of modified SSC 5X solution (750 mM NaCl, 75 mM sodium citrate, 0.05 v% sodium dodecyl sulfate, and 0.05 wt% lauroylsarcosine). Wells were blocked with 3% wt BSA PBS buffer pH 7.2 for 2 hours at 4°C under stirring and washed once with 200 µL of SSC 5X solution. Then 100 µL at 16.5 µM of capture oligonucleotide DIG-5' (5‘-GACCCCAAAATCAGCGAAAT) were added and the plates were kept 2 hours at 4°C under stirring. In parallel, 60 µL of the target DNA at 43 nM were denatured for each concentration at ambient temperature during 15 minutes in a 0.1 M NaOH, 0.01% Tween 20 solution. Then, 540 µL of detection oligonucleotide 3'-biotin (5'-CAGATTCAACTGGCAGTAACCAGA) at 40 nM in modified SSC 5X solution were added. After washing the wells with 200 µL of modified SSC 5X solution, 100 µL of this target DNA / detection oligonucleotide conjugate solution were added and the plates were conserved at 40°C under stirring during 30 minutes. Last, 100 µL of $YVO_4:Eu^{3+}$ nanoparticle-streptavidin conjugates at 4.5 nM were added and the samples were incubated for 1 hour at 4°C under stirring. The wells were then washed 5 times with modified SSC 5X solution and 2 times with PBS buffer prior to readout with our homemade UV microplate reader.

**PCR**

PCR amplification was performed on the plasmid pEX-A128-nCOV_N1 (Eurofins) using the forward primer 5'-GACCCCAAAATCAGCGAAAT and the reverse primer 5'-TCTGGTTACTGCCAGTTGAATCTG (Eurogentec). The PCR mix was performed using the Q5 high fidelity PCR kit (Biolabs) following manufacturer recommendations with a hybridization temperature of 50°C and an elongation step of 2 minutes. The PCR product of 72 bp was purified using the GeneJet PCR purification kit (K0701, Thermofisher) following manufacturer's instructions. The result was analyzed using electrophoresis on an agarose gel revealed by BET (ref. SM1213, Thermofisher), imaged using the Chemidoc MP imaging system (Biorad) and the analysis of the gel image was performed using ImageJ (Fig. 3a).
